# Plasmid DNA analysis of pristine groundwater microbial communities reveal extensive presence of metal resistance genes

**DOI:** 10.1101/113860

**Authors:** Ankita Kothari, Yu-Wei Wu, Marimikel Charrier, Lara Rajeev, Andrea M. Rocha, Charles J. Paradis, Terry C. Hazen, Steven W. Singer, Aindrila Mukhopadhyay

## Abstract

Native plasmids constitute a major category of extrachromosomal DNA elements responsible for harboring and transferring genes important in survival and fitness. A focused evaluation of plasmidomes can reveal unique adaptations required by microbial communities. We examined the plasmid DNA from two pristine wells at the Oak Ridge Field Research Center. Using a cultivation-free method that targets plasmid DNA, a total of 42,440 and 32,232 (including 67 and 548 complete circular units) scaffolds > 2 kb were obtained from the two wells. The taxonomic distribution of bacteria in the two wells showed greater similarity based on their plasmidome sequence, relative to 16S rRNA sequence comparison. This similarity is also evident in the plasmid encoded functional genes. Among functionally annotated genes, candidates providing resistance to copper, zinc, cadmium, arsenic, and mercury were particularly abundant and common to the plasmidome of both wells. The primary function encoded by the most abundant circularized plasmid, common to both wells, was mercury resistance, even though the current ground water does not contain detectable levels of mercury. This study reveals that the plasmidome can have a unique ecological role in maintaining the latent capacity of a microbiome enabling rapid adaptation to environmental stresses.

## Introduction

Sudden changes in environmental stress can be better dealt with by horizontal gene transfer (HGT) and gene duplication events than by slower mutation and selection mechanisms (Aminov, 2011). Plasmids are important in HGT and are critical in facilitating genome restructuring by providing a mechanism for distributing genes that provide a selective advantage to their host. Typically, plasmids have a modular structure, containing several functional genetic modules. Plasmids typically vary from 5-500 kb in size, although plasmids as small as 2 kb (Biet *et al.*, 2002; Kobori *et al.*, 1984; Rozhon *et al.*, 2010) to as large as more than 1 Mb in size (Harrison *et al.*, 2010; Finan *et al.*, 1986) have been reported. Despite their importance, plasmids are rarely examined by themselves and only a handful of studies have focused on extrachromosomal DNA in microbial communities (Kav *et al.*, 2012; 2013; Mizrahi, 2012; Jørgensen *et al.*, 2014a; Zhang *et al.*, 2011).

Due to the role of plasmids in environmental stress adaptation, they may be particularly important in sites that have historical significance in accumulated contaminants. The Oak Ridge Field Research Center (OR-FRC) site at the Y-12 Federal Security Complex in Oak Ridge, TN, is known to be contaminated with radionuclides (e.g., uranium and technetium), heavy metals, nitrate, sulfide, and volatile organic compounds. This is a well-characterized experimental field site and has been used for studying the environmental impacts of contaminants in the past (Watson *et al.*, 2004; Hemme *et al.*, 2010). To better understand the most prevalent functionality that is horizontally transferred and how the memory of exposure to contaminants persists in the environment we studied the plasmidome of the pristine wells, located in the same watershed as the contaminated wells, but 6 miles from the contaminated areas of the watershed (Smith et al., 2015).

A plasmidome has been described to be the entire plasmid content in a given environment that is resolved by metagenomic approaches during high-throughput-sequencing experiments (Dib *et al.*, 2015). Plasmidome analyses have been performed in cow rumen (Kav *et al.*, 2012; Mizrahi, 2012), rat caecum (Jørgensen *et al.*, 2014b), and industrial wastes (Zhang *et al.*, 2011), all of which are highly abundant in bacteria. In contrast, the environmental stress at OR-FRC has resulted in a substantial decrease in cells counts, and species diversity (Hemme *et al.*, 2016), presenting unique challenges in plasmid isolation. Examination of plasmid DNA from this site also poses challenges due to the range in size and diversity of the plasmid DNA expected to arise from the fluctuating microbial community of this highly uncontrolled/variable environment. We present an optimized method that overcomes several of these challenges, and the first study to selectively isolate and analyze the plasmidome from ground water samples at the OR-FRC. We provide insights into the composition and structure of the plasmidome of two pristine wells, GW456 and GW460 in context of corresponding geochemical and metagenome data. We present several novel findings from the plasmidome analysis, including the presence of common scaffolds in the two wells, and an abundance of plasmid-encoded metal resistance genes.

## Materials and Methods

### Optimization of plasmid DNA isolation methods using a model system

The model system of a 1:1:1 mixture of *Desulfovibrio vulgaris* Hildenborough (ATCC 29579) containing a 202 kb native plasmid (pDV1), *Escherichia coli* DH1 (ATCC 33849) containing a 48 kb fosmid (fSCF#19) (Ruegg *et al.*, 2014), and the *E. coli* strain J-2561 containing a 5 kb (pBbS5c) plasmid was prepared using cells grown to OD 1. *Desulfovibrio* was grown in LS4D supplemented with 0.1% (w/v) yeast extract (Ray *et al.*, 2014) while *E. coli* was grown in LB media. This mixture was serially diluted ten-fold, stored at −80 °C and used to test, compare and optimize plasmid detection via quantitative polymerase chain reaction (qPCR). Two alkaline hydrolysis methods were compared to preferentially isolate plasmid DNA: Birnboim and Doly, 1979 and Anderson and Mckay, 1983. Residual linear chromosomal DNA fragments were minimized by Plasmid-Safe ATP-Dependent DNase (Epicentre, Madison, WI, USA) treatment for 24 - 48 h at 37 °C. The presence of genomic DNA was tested by PCR using 16S rRNA universal primers (BAC338F: 5’-ACTCCTACGGGAGGCAG-3’ and BAC805R: 5’-GACTACCAGGGTATCTAATCC-3’) (Stevenson and Weimer, 2007). If 16S rRNA PCR product was visible on a 1 % agarose gel, another overnight digestion reaction was performed until the product could no longer be visualized. The DNase was inactivated at 70 °C for 30 min. The DNA was then amplified with phi29 DNA polymerase (New England Biolabs, Ipswich, MA, USA) (Kav *et al.*, 2012) at 4, 18 or 30 °C for 168, 25, and 24 h respectively. Plasmid isolation was checked via qPCR against a specific plasmid-borne gene on all three plasmids. qPCR was performed using the SsoAdvanced Universal SYBR Green Supermix (Bio-Rad, Hercules, CA, USA) as per the manufacturers protocol. Total DNA from *D. vulgaris* Hildenborough was used as a control for the 202 kb primers and the plasmid DNA coding for pBbS5c was used as a control for the 5 kb primers.

### Sample collection

Water samples were collected from two uncontaminated deep groundwater wells of the Department of Energy’s OR-FRC, Tennessee (Dougherty *et al.*, 2014) - GW456 (11th November 2014; GPS coordinates 35°56’28.4”N 84°20’10.3”W) and GW460 (1^st^ December 2014; GPS coordinates 35°56’27.9”N 84°20’10.7”W) (Supplementary Information 1). These wells are about 60 feet away from each other. Prior to collection of samples, approximately 5-20 l of groundwater was pumped until temperature, pH, conductivity, and oxidation-reduction (redox) values were stabilized to purge the well and the line of standing water. An additional 25 ml groundwater, along with 25 ml glycerol was stored at −80 °C for any further required analyses. The water was collected with a peristaltic pump using low-flow to minimize drawdown of the well. Bulk water measurements and geochemical sample collections (Smith *et al.*, 2015) were conducted. For 16S rRNA analysis and plasmid isolation a total of 8 and 5 l of water, respectively, was filtered through a 144 mm diameter 10 μm Nylon filter (Sterlitech Corporation, Kent, WA, USA). Filters were immediately stored on dry ice in 50 ml falcon tubes until transported to the −80 °C freezer.

### Geochemical Measurements

Temperature, pH, conductivity, redox, and dissolved oxygen were measured at the wellhead using an In-Situ Troll 9500 (In-situ Inc., Fort Collins, CO, USA). Sulfide and ferrous ion groundwater concentrations were determined using the USEPA Methylene Blue Method (Hach 8131) and 1,10-Phenanthroline Method (Hach 8146), respectively, and analyzed with a field spectrophotometer (Hach DR 2800). All other biological and geochemical parameters were measured as previously described (Smith *et al.*, 2015). Mercury analysis was performed on samples containing 25 ml groundwater and 25 ml glycerol by Oxidation, Purge, Trap, and Cold Vapor Atomic Fluorescence Spectrometry −1631E at ALS Environmental, Kelso, WA, USA.

### Plasmid isolation from environmental samples

The filters containing cells from the wells GW456 and GW460 were thawed to room temperature. The filter was cut into tiny pieces in a sterile petri dish using sterilized forceps and scissors and split into two 50 ml falcon tubes. About 1.33 × 10^5^ cells, of each plasmid-containing control strain, were added to each tube. The plasmid isolation method (Anderson and Mckay, 1983) was used with the following modifications. The volumes of all reagents were multiplied twenty times to immerse each half filter. Before the addition of lysozyme (Sigma-Aldrich, St. Louis, MO, USA), the samples were heated to 37 °C with gentle inversion for 10 mins and vortexed with 0.1 mm Disrupter Beads (Scientific Industries, Bohemia, NY, USA) at medium setting for 5 mins. After the addition of sodium chloride, the liquid was transferred into 50 ml Phase Lock Gel Heavy tubes (5-Prime). 14.5 ml of 25:24:1 phenol:chloroform:isoamyl alcohol was added to each tube, thoroughly mixed and centrifuged for 5 mins at 1500 g (Beckman Coulter Allegra 25R centrifuge). The upper phase was transferred to a fresh phase lock tube. 14.5 ml of 24:1 chloroform-isoamyl alcohol was added and centrifuged for 5 mins at 1500 g. The upper phase was transferred to a 50 ml falcon tube and precipitated with an equal volume of isopropanol. The extractions from each half of the filter were recombined, incubated on ice for 1 h, followed by centrifugation for 5 mins at 8000 g. The excess isopropanol was removed and the pellet was resuspended in 1 ml 10 mM Tris, 1 mM EDTA, pH 7, and transferred to a 1.6 ml tube and dehydrated down to 50 μl with Vacufuge plus (Eppendorf, V-AQ, 45 °C). The remnant linear DNA fragments were removed by Plasmid-Safe ATP-Dependent DNase (37 °C for 48 h with double the recommended ATP and enzyme amount) and the lack of genomic DNA contamination was confirmed by PCR with degenerate 16S rRNA primers (Supplementary Information 2). The plasmid DNA was amplified with phi29 DNA Polymerase for 6 days at 18 °C. This was followed by ethanol precipitation and nanodrop to concentrate and quantify the DNA.

### Plasmid sequencing and bioinformatics

The plasmid DNA was sequenced at the Vincent J. Coates Genomics Sequencing Laboratory at UC Berkeley using the Illumina MiSeq reagent v3 kit (paired-end protocol). As reported previously (Jørgensen *et al.*, 2014b), IDBA-UD (Peng *et al.*, 2012) was used for *de novo* read assembly with the parameter “--pre_correction”. Assembled sequences were searched against the SILVA 16S rRNA database (Quast *et al.*, 2013) using BLASTN; all scaffolds with > 200 bp identity to 16S rRNA were removed from further analysis. To exclude the control plasmids, all sequences with more than 95 % identity to these plasmids (minimum alignment length 1000 bp) were also removed. The taxonomic and functional composition of the dataset was analyzed with the MG-RAST server (Glass and Meyer, 2011) using similarity to the SEED database (with a maximum E-value of ≤10^−5^) (Overbeek *et al.*, 2005).

We modified a pipeline method for post-assembly detection of circularity among scaffolds (Jørgensen *et al.*, 2014b) with the following criteria to identify the complete closed circular scaffolds “circular_scaffolds”: a) Scaffold length >2 kb; b) >34 bp homology (e-value > 1e-5) at the ends of the scaffold in the correct direction; c) At least two read pairs mapped on opposite ends of the contig, a maximum of 500 bp from the end. The complete pipeline with Perl scripts can be found at https://github.com/yuwwu/detect-circ-plasmid. The “circular_scaffolds” were subjected to annotation using components from the RAST (Rapid Annotations using Subsystems Technology) toolkit (RASTtk) with the Department of Energy Systems Biology Knowledgebase, KBase (http://kbase.us).

The obtained “all_scaffolds” and “circular_scaffolds” plasmid databases were compared with 1) A CLAssification of Mobile genetic Elements (ACLAME) (Leplae *et al.*, 2004; Lima-Mendez *et al.*, 2010), 2) antiBacterial biocide and Metal resistance genes database (BacMet)(Pal *et al.*, 2014), 3) Toxin Antitoxin DataBase (TADB) (Shao *et al.*, 2011) 4) Antibiotic Resistance genes DataBase (ARDB) (Liu and Pop, 2009) and 5) Comprehensive Antibiotic Resistance Database (CARD) (McArthur *et al.*, 2013) databases. The analyses were performed as follows: 1) ACLAME: The ACLAME plasmid proteins and MGE (Mobile Genetic Elements) families were downloaded from the ACLAME website. The plasmid genes from both wells were mapped against the plasmid proteins using BLAST with e-value cutoff 1e-3. The BLAST tabulated results were parsed to obtain the taxonomic distributions of the plasmid genes by mapping the BLAST results to the MGE families, which consists of the taxonomic information. 2) BacMet: The perl script BacMet-Scan.pl version 1.1, the predicted resistance genes datasets and the experimentally confirmed resistance genes dataset was downloaded from the BacMet website. The BacMet-Scan.pl was executed using default parameters (-blast -e 1 -l 30 -p 90) to generate the tabulated report against both predicted and experimentally-confirmed datasets. 3) TADB: The database was downloaded from the TADB website version 1.1 (http://202.120.12.135/TADB2/) followed by BLAST with the following parameters: -evalue 1e-3 -min_target_seqs 1. 4) ARDB: The perl script ardbAnno.pl and ardbAnno.pm were downloaded from the ARDB website along with the resistance gene dataset. The plasmid genes from both wells were mapped against the resistance gene dataset using the scripts with default parameters. 5) CARD: CARD and software RGI (Resistant Gene Identifier) databases were downloaded from the CARD website (https://card.mcmaster.ca/home). The script rgi.py was used to search the predicted plasmid genes against the CARD database with default parameters followed by parsing using a customized perl script.

### 16S rRNA sequencing

Genomic DNA was extracted using the modified Miller DNA extraction method (Smith *et al.*, 2015) followed by purification and concentration using a Genomic DNA Clean & Concentrator Kit (Zymo Research, Irvine, CA). DNA quality was determined by the NanoDrop spectrophotometer (Thermo Scientific, Waltham, MA) and its concentration was determined by a Qubit 2.0 Fluorometer (Life Technologies, Carlsbad, CA). The V4 region, of both bacterial and archaeal 16S rRNA genes, was amplified using a two-step PCR approach. The primers [515F, 5′-GTGCCAGCMGCCGCGGTAA-3′ and 806R, 5′-GGACTACHVGGGTWTCTAAT-3′ were used without added sequencing components in the first step to avoid additional bias. To increase the base diversity in sequences of sample libraries, phasing primers were used in the second step PCR. Spacers of different length (0-7 bases) were added before the forward and reverse primers, which shifts sequencing phases amongst different community samples from both directions. Sequencing was performed on the Illumina MiSeq platform (Smith *et al.*, 2015).

The resulting 16S rRNA sequence data were processed using custom python scripts (https://github.com/almlab/SmileTrain) that call USEARCH for quality filtering and overlapping paired end reads, and biopython (Cock *et al.*, 2009) for file format input and output. The sequences were then progressively clustered to 90 % with UCLUST (Edgar, 2010), aligned to the SILVA database with mothur, align.seqs and processed with distribution-based clustering (DBC) as previously described (Preheim *et al.*, 2013) with k_fold 10 to remove sequencing errors. OTU representatives were defined during clustering as the most abundant sequences in the OTU. Chimeras were identified with UCHIME (Edgar *et al.*, 2011) and removed. Taxonomic identification was performed with RDP (Wang *et al.*, 2007) using 0.50 as a confidence threshold for taxonomic classification at every level. The OTU table data was then converted to a biom format to analyze diversity and taxa summaries in Qiime.

## Results and Discussion

To capture plasmids of different sizes, present in unpredictable copy numbers, we optimized a plasmid DNA isolation method applicable to environmental samples. For this we used a model system comprising of three bacterial strains harboring plasmids of sizes 5 kb, 48 kb and 202 kb. We isolated the plasmids with two alkaline hydrolysis methods, followed by Plasmid-Safe ATP-Dependent DNAse digest to remove linear DNA fragments, and phi29 amplification of the isolated plasmid DNA (Fig 1). We compared two plasmid isolation methods (Birnboim and Doly, 1979; Anderson and Mckay, 1983) and three phi29 incubation temperatures 4, 18 and 30 °C at two different cell concentrations (5 × 10^5^ or 5 × 10^4^ cells per strain). We used qPCR with a plasmid-borne gene on each plasmid to determine the optimal condition for detecting all three plasmids.

We found that that smaller the size of the plasmid, the easier it was to isolate and detect (Supplementary Information 3a). This was expected since smaller plasmids are typically maintained in high copy numbers and smaller ssDNA fragments re-anneal more efficiently during alkaline hydrolysis, increasing the chances of their isolation. Of the parameters tested, the optimal plasmid isolation conditions to detect all three plasmid sizes was to use the Anderson and Mckay, 1983 alkaline hydrolysis method with phi29 amplification at 18 °C. Decreasing the number of cells of each strain 10-fold resulted in detection of only the 5 kb plasmid (data not shown). Since the ground water samples from the OR-FRC were present on a filter paper, we evaluated the efficiency of our optimized plasmid isolation method using model system strains on identical filters. The qPCR based analysis revealed that the filter did not interfere with the extraction procedure (Supplementary Information 3b) and the extraction could be improved by cutting the filter into smaller pieces and vortexing with beads. The optimal number of cells remained in the range of 5 × 10^5^ cells for the detection of all the three plasmids.

**Fig 1.**
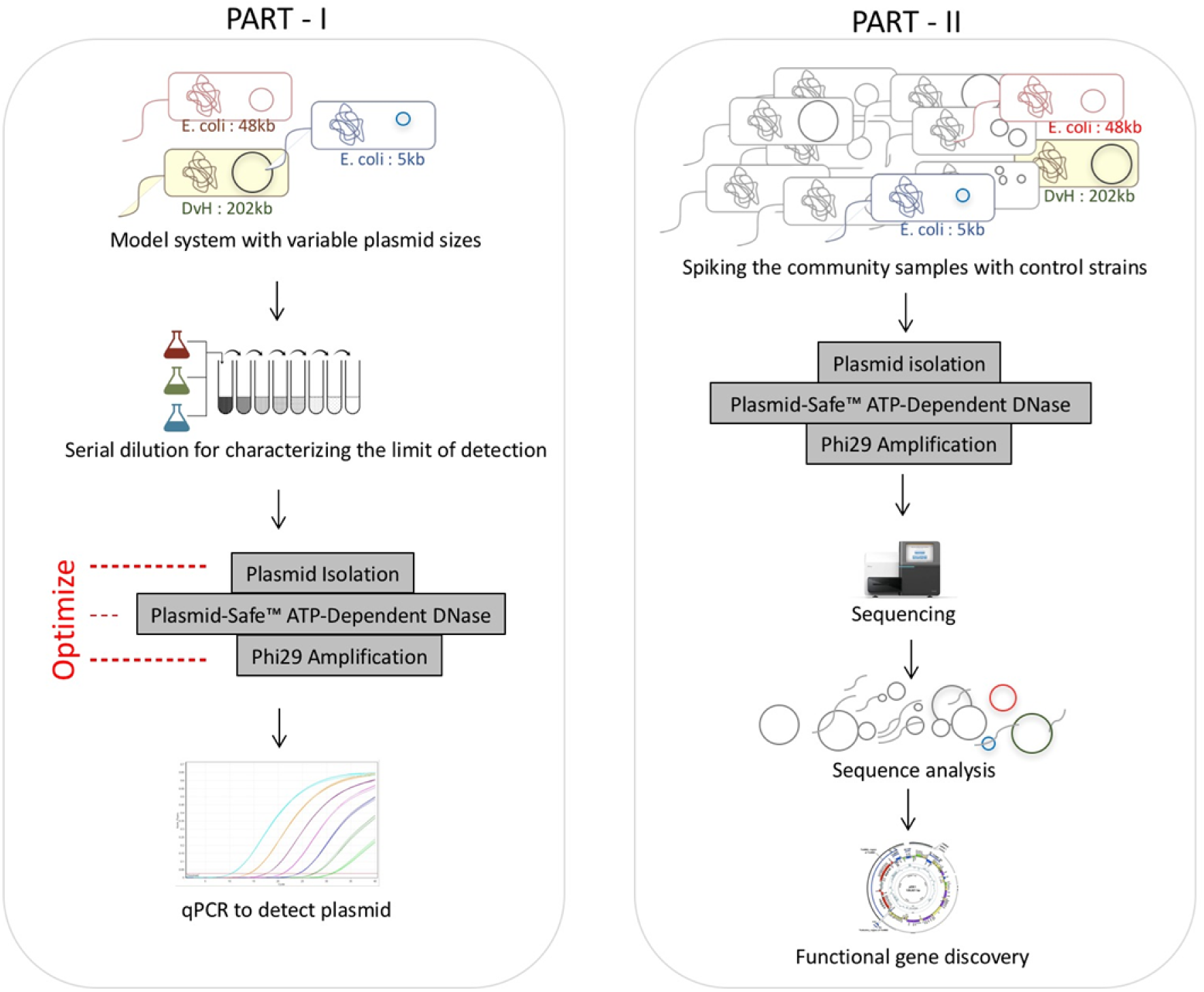
An overview of the project work flow. Part I involved optimizing plasmid isolation methods using a model system containing two *E. coli* strains and one *Desulfovibrio vulgaris Hildenborough* (DvH) with 5, 48 and 202 kb plasmid sizes. Successful isolation of plasmid was confirmed by qPCR against a specific plasmid borne gene. Part II involved using the optimized method to isolate plasmids from the environmental sample, followed by sequence analysis.

The optimized method allows us to detect plasmid DNA spanning 5-200 kb via qPCR from cells present at 5 × 10^5^ or greater, from filtered ground water. We used the optimized method on samples from two pristine wells of the OR-FRC, GW456 and GW460 followed by sequencing (for increase the sensitivity of plasmid detection). The resulting plasmidome was sequenced and generated datasets with 34 and 76 million reads, respectively (Table 1). In each case, the reads were trimmed and any scaffolds associated with 16S rRNA sequences or the spiked control plasmids were removed. Of the spiked control plasmids, only the 5 and 48 kb plasmids were detected, perhaps because 1) it is easier to detect small plasmid sizes 2) the number of native plasmids in the environmental sample was overwhelming. We observed a maximum scaffold size of 1.16 Mb in GW456 and 0.63 Mb in GW460. Based on the earlier reports for plasmid sizes (Harrison *et al.*, 2010), they are well within the expected size range for plasmids. Overall, the wells GW456 and GW460 generated 462,471 and 521,975 scaffolds with a mean scaffold length of 2059 bp and 1739 bp respectively. This is higher than the 469 bp from earlier reports (Kav *et al.*, 2013), and indicate that we were able to capture higher plasmid diversity. Interestingly, 1188 scaffolds show more than 90 % identity and more than 90 % query coverage between the two wells. The common scaffolds range from size 2.0 - 175.9 kb, with an average size of 8.5 kb.

**Table 1.**
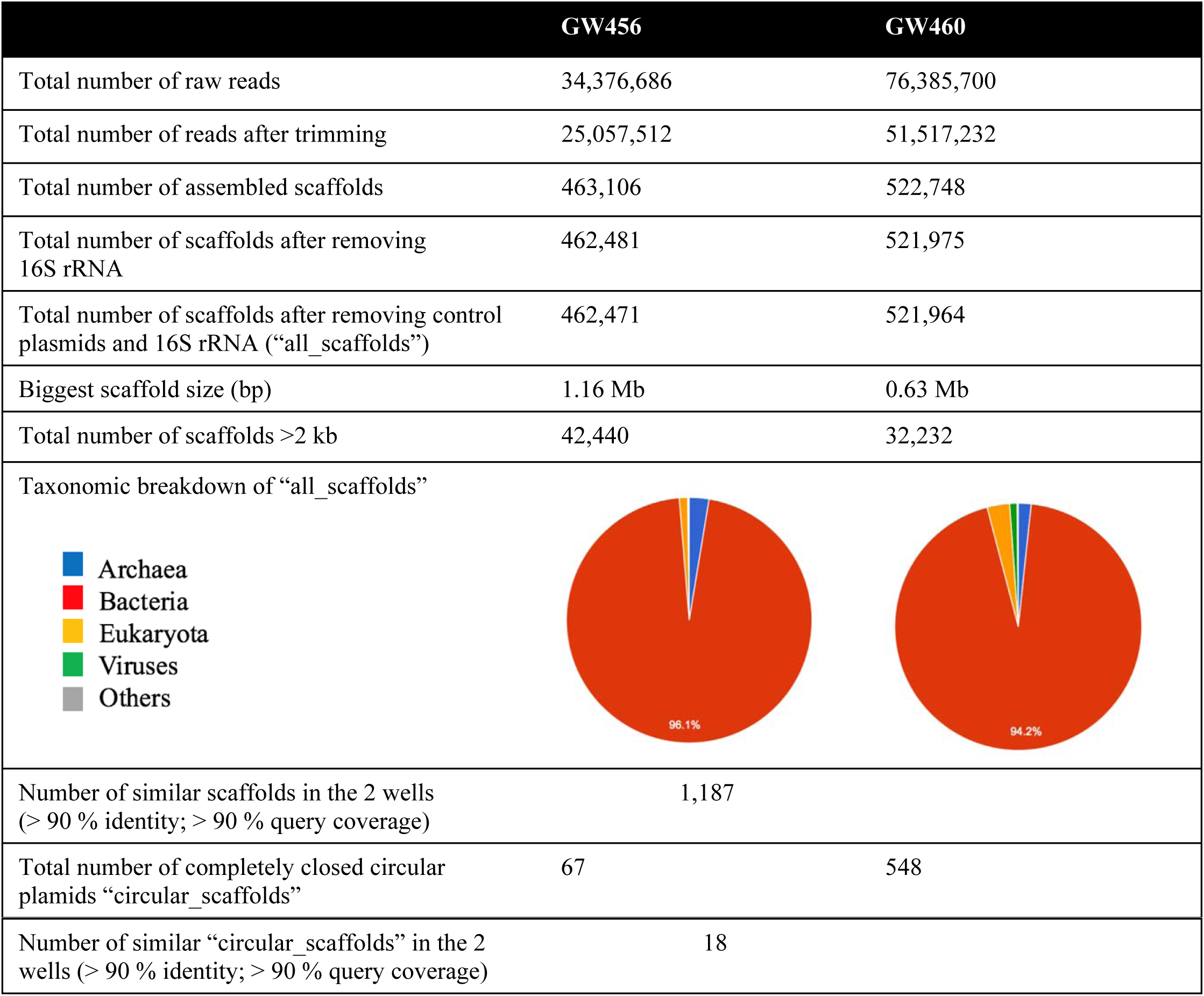
Plasmidome sequencing analysis from the uncontaminated wells GW456 and GW460 in Oak Ridge – Field Research Center.

## Taxonomic distribution of the plasmidomes show striking similarities between the two wells

For each well, the taxonomic distributions based on 16S rRNA and that of the plasmidome revealed the presence of similar taxa, albeit the abundance differs (Fig 2). The wells at the OR-FRC are known to exhibit daily fluctuations in the 16S rRNA distribution. Correspondingly, comparison of the 16S rRNA profiles between the two wells showed considerable differences in the distribution of phyla. In contrast, the taxonomic distributions of the plasmidome of the two wells were significantly more similar to each other (Fig 2). Thus, the plasmidomes of the two wells are more like each other despite the difference in 16S rRNA profiles.

**Fig 2.**
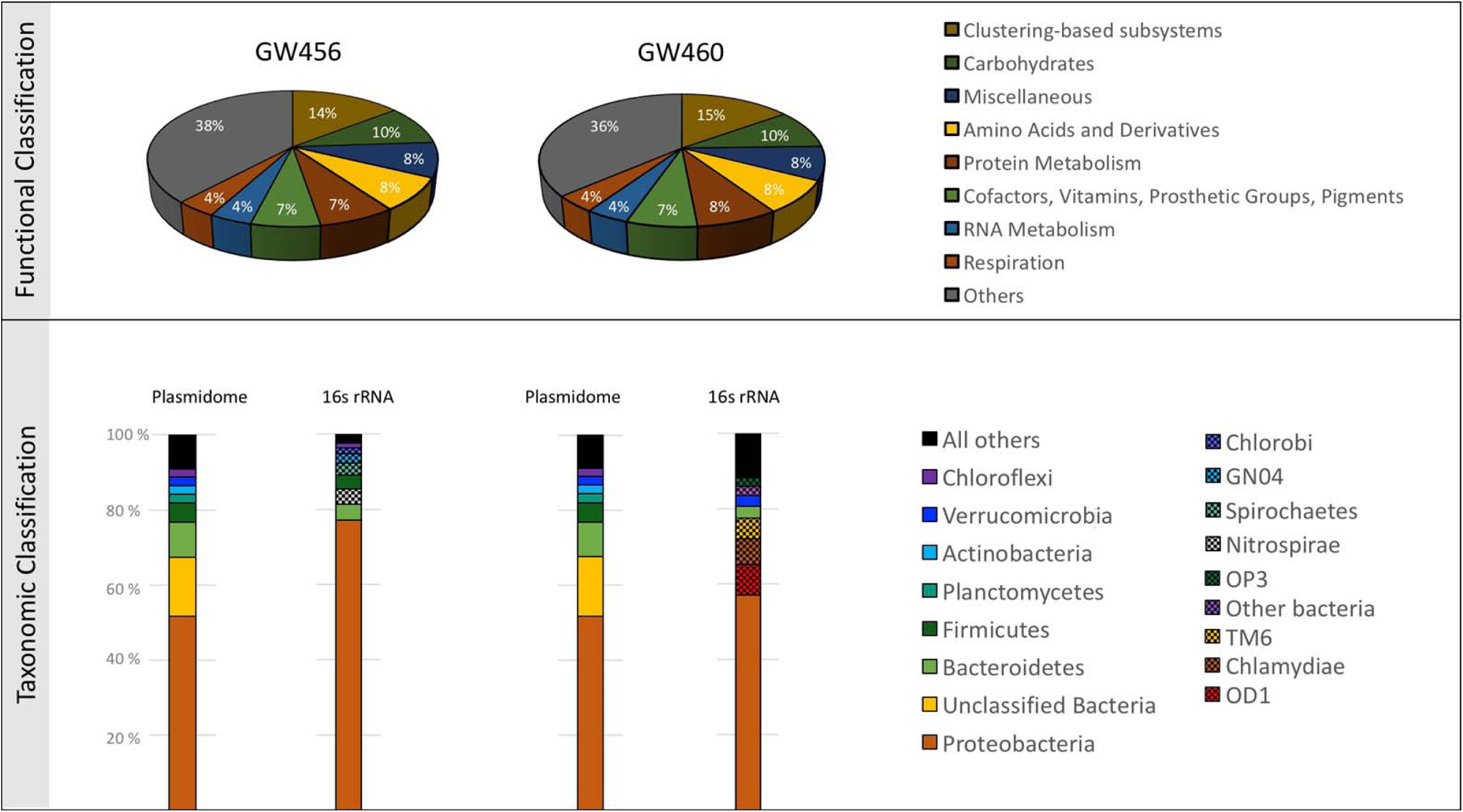
a) Functional classification of “all_scaffolds” into SEED subsystem categories using MG-Rast. b) Taxonomic classification of “all_ scaffolds” into phyla based on 16S rRNA (QIIME) and plasmidome (MG-Rast) sequences. The MG-Rast analysis was based on the lowest common ancestor (Parameters: 1e-5 maximum e-value cut off, 60 % maximum identity cut off, 15 bp minimum alignment length cut off).

Phylogenetic assignment performed by the SEED subsystems database showed that most of the scaffolds (> 94 %) were assigned to Bacteria, with minor representation of the Archaea, Eukaryota and Viruses domains (Table 1). At the domain, phyla and order levels, the taxonomic distribution of the sequenced scaffolds from both the wells were similar. The most dominant phylum (in 16S rRNA and plasmidome profiles) was Proteobacteria (Fig 2). Within this phylum, the classes Alpha-Beta-, Gamma- and Delta-proteobacteria were highly abundant in both wells. A previous study on the OR-FRC has shown that Proteobacteria such as *Burkholderia* and *Pseudomonas* were the most abundant lineages in the pristine wells while *Rhodanobacter* was dominant in the contaminated wells (Hemme *et al.*, 2015). In this study we observed a high abundance of Proteobacteria, however none of the genes encoded on plasmids were annotated to belong to *Rhodanobacter.*

One explanation for the unexpected plasmidome similarity between the wells could be the geographical proximity of the wells that allows the flowing water to be shared. Alternately, it may be that that there are limited variations in the genetic modules that constitute a plasmid. We speculate that the former is the more likely explanation although the second cannot be ruled out given that the modules on the plasmids (e.g. scaffold_5343 and scaffold_67) show very high similarity to other plasmids reported from diverse geographical locations across the globe (see below).

## Plasmid-borne functionally annotated genes are similar in the two wells

Functional classification of the scaffolds into SEED subsystems categories revealed a highly similar distribution of plasmid-borne genes between the two wells (Fig 2a). The most highly represented subsystems were “Carbohydrates”, “Amino Acids metabolism”, and “Clustering based sub-systems”. Similar categories were known to be abundant in the plasmidome of rumen bacteria (Kav *et al.*, 2013). One of the highly abundant categories in SEED level-4 classification, was “Resistance to antibiotics and toxic compounds”. In this category, the cobalt, zinc and cadmium resistance genes were the most abundant. Genes involved in iron transport were also highly abundant in both wells. Functional classification of the two wells based on Clusters of Orthologous Groups (COG) was also similar (Supplementary Information 4). The only category significantly different between the two wells was the presence of phage capsid proteins in GW460.

We successfully circularized 67 and 548 of the sequenced scaffolds from GW456 and GW460 respectively. The plasmid size distribution of “all_scaffolds” and “circular_scaffolds” of the two wells were similar (Supplementary Information 5). The average size of “circular_scaffolds” was 19 kb and 4.5 kb from the wells GW456 and GW460, respectively. The GC content distribution of the genes from both wells for “all_scaffolds” and “circular_scaffolds” revealed a bias towards high GC content in the well GW460 (Supplementary Information 6). The abundant circular plasmids from both wells encoded metal resistance genes, indicating their significance in this environment (Table 2a, b). Consistent with the overall taxonomic similarity in the sequenced scaffolds from the two wells, 18 circular plasmids were also common to both wells, with >99.8 % sequence identity and >93.6 % query coverage (Table 2c). Some of these closed plasmids were associated with known *rep* and *mob* genes involved in plasmid replication and mobilization. Certain others did not encode any known phenotypic traits, and were hence cryptic (Novick *et al.*, 1976). The latter could potentially serve as an important source for discovery of novel functional genes and replication systems (Jørgensen *et al.*, 2014b). Based on the genes associated with plasmids, the “circular_scaffolds” were annotated as conjugative, mobilizable, or non-mobilizable (Smillie *et al.*, 2010) plasmids (Fig 3). As observed previously (Smillie *et al.*, 2010), the non-mobilizable plasmids were highly dominant in both wells.

**Table 2a.** The top twenty most abundant “circular_scaffolds” from well GW456

| GW456 plasmid | Coverage | Plasmid size (kb) | Resistance Genes | Plasmid associated genes |
| --- | --- | --- | --- | --- |
| scaffold_5343 | 869 | 7993 | Metal Resistance (Mercury) | Mobilization protein-coding gene *mobA*, *mobC*,<br>Replication protein-coding gene *repA* |
| scaffold_2832 | 445 | 12888 | Metal Resistance (Cobalt Zinc, Cadmium) | Plasmid partition protein-coding gene *parA*,<br>Antitoxin-coding gene *higA*,<br>Replication protein-coding gene *repA*,<br>Mobilization protein-coding gene *mobA* |
| scaffold_6029 | 288 | 7364 | - | Mobilization protein-coding gene |
| scaffold_15459 | 248 | 3832 | - | Mobilization protein-coding gene *mobA* |
| scaffold_11977 | 196 | 4554 | - | Doc Toxin-coding gene |
| scaffold_6372 | 186 | 7077 | - | - |
| scaffold_215 | 139 | 96159 | - | Antitoxin-coding gene *higA*,<br>Mobilization protein-coding gene *mobA*,<br>RelE/StbE replicon stabilization toxin-coding gene,<br>RelB/StbD replicon stabilization protein-coding gene (antitoxin to RelE/StbE),<br>IncF plasmid conjugative transfer pilus assembly protein-coding genes *traH*, *traC*, *traC*, *traB*<br>Plasmid partition protein-coding gene *parA* |
| scaffold_1417 | 105 | 21696 | - | Replication protein-coding gene *repA*,<br>Plasmid partition protein-coding gene *parA*,<br>Toxin-coding gene *higB*,<br>plasmid maintenance system antidote protein - XRE family,<br>Coupling protein-coding gene *virD4* |
| scaffold_9451 | 102 | 5372 | - | - |
| scaffold_117 | 84 | 287339 | Phage resistance protein | - |
| scaffold_24704 | 74 | 2796 | Metal Resistance (Zinc) | - |
| scaffold_18901 | 73 | 3916 | - | - |
| scaffold_12628 | 35 | 4388 | - | - |
| scaffold_35161 | 29 | 2218 | - | - |
| scaffold_12673 | 28 | 4335 | No | Toxin-coding gene *yoeB*<br>Antitoxin-coding gene *yefM* |
| scaffold_130 | 19 | 94434 | Metal Resistance (Cobalt Zinc,<br>Cadmium, Copper, Arsenic),<br>Antibiotic Resistance<br>(Fosfomycin,<br>Spectinomycin),<br>Metal uptake (Potassium<br>uptake) | - |
| scaffold_3845 | 17 | 10328 | - | - |
| scaffold_34293 | 10 | 2253 | - | - |
| scaffold_23147 | 8 | 2916 | - | - |
| scaffold_31933 | 7 | 2363 | - | - |
| scaffold_27134 | 6 | 2632 | - | - |

**Table 2b:** The top twenty most abundant “circular_scaffolds” from well GW460

| GW460 plasmid | Coverage | Plasmid size (kb) | Resistance Genes | Plasmid associated genes |
| --- | --- | --- | --- | --- |
| scaffold_23986 | 2593 | 2300 | - | - |
| scaffold_28338 | 1799 | 2059 | - | - |
| scaffold_8056 | 910 | 4782 | - | - |
| scaffold_19698 | 890 | 6427 | - | - |
| scaffold_10032 | 617 | 7993 | Metal Resistance (Mercury) | Mobilization protein-coding genes *mobA*, *mobC*,<br>Replication protein-coding gene *repA* |
| scaffold_10418 | 542 | 4017 | - | RelB/StbD replicon stabilization protein-coding gene (antitoxin to RelE/StbE),<br>RelE/StbE replicon stabilization toxin-coding gene |
| scaffold_15696 | 424 | 3049 | - | - |
| scaffold_13367 | 379 | 3388 | - | - |
| scaffold_7597 | 357 | 9688 | - | - |
| scaffold_28387 | 325 | 3015 | - | - |
| scaffold_19474 | 325 | 2632 | - | Replication protein-coding gene |
| scaffold_5179 | 312 | 6544 | - | Mobilization protein-coding genes *mobC* |
| scaffold_24781 | 298 | 2253 | - | - |
| scaffold_25039 | 252 | 2238 | - | - |
| scaffold_8800 | 210 | 4498 | - | - |
| scaffold_16750 | 191 | 2916 | - | - |
| scaffold_2324 | 171 | 12888 | Metal Resistance (Cobalt Zinc, Cadmium, Mercury) | Plasmid partition protein-coding gene *parA*,<br>Replication protein-coding gene *repA*, |
|  |  |  |  | Mobilization protein-coding genes *mobA*,<br>Coupling protein-coding gene *virD4* |
| scaffold_23312 | 154 | 5574 | - | Mobile element protein,<br>Integrase |
| scaffold_3529 | 149 | 8684 | - | Coupling protein-coding gene *virD4*,<br>Mobilization protein-coding genes *mobA* |

**Table 2c:** Eighteen circular plasmids that displayed >99.8 % sequence identity with >93.6 % query coverage in the two wells. Their scaffold number, coverage, plasmid size, resistance gene(s) and the plasmid associated gene(s) encoded are tabulated

| GW456 plasmid | Coverage | GW460 plasmid | Coverage | Mobility | Plasmid size (bp) | Resistance Genes | Plasmid associated genes |
| --- | --- | --- | --- | --- | --- | --- | --- |
| scaffold_5343 | 869 | scaffold_10032 | 617 | Mobilizable | 7993 | Metal Resistance (Mercury) | Mobilization protein-coding gene *mobA*,<br>*mobC*,<br>Replication protein-coding gene *repA* |
| scaffold_2832 | 445 | scaffold_2324 | 171 | Conjugative | 12888 | Metal Resistance (Cobalt Zinc, Cadmium) | Plasmid partition protein-coding gene *parA*,<br>Antitoxin-coding gene *higA*,<br>Replication protein-coding gene *repA*,<br>Mobilization protein-coding gene *mobA* |
| scaffold_11977 | 196 | scaffold_8648 | 64 | Non-Mobilizable | 4554 | - | Doc Toxin-coding gene |
| scaffold_215 | 139 | scaffold_45 | 79 | Conjugative | 96159 | - | Antitoxin-coding gene *higA*,<br>Mobilization protein-coding gene *mobA*,<br>RelE/StbE replicon stabilization toxin,<br>RelB/StbD replicon stabilization protein-coding gene (antitoxin to RelE/StbE),<br>IncF plasmid conjugative transfer pilus assembly protein-coding genes *traH*,<br>*traC*, *traC*, *traB*,<br>Plasmid partition protein-coding gene *parA* |
| scaffold_1417 | 105 | scaffold_1021 | 11 | Conjugative | 21696 | - | Replication protein-coding gene *repA*,<br>Plasmid partition protein-coding gene *parA*,<br>Toxin-coding gene *higB*,<br>plasmid maintenance system antidote protein - XRE family, |
|  |  |  |  |  |  |  | Coupling protein-coding gene *virD4* |
| scaffold_9451 | 102 | scaffold_6875 | 63 | Non-Mobilizable | 5372 | - | - |
| scaffold_18449 | 83 | scaffold_13367 | 379 | Non-Mobilizable | 3388 | - | - |
| scaffold_24704 | 74 | scaffold_17814 | 52 | Non-Mobilizable | 2796 | Metal Resistance (Zinc) | - |
| scaffold_18901 | 73 | scaffold_13695 | 25 | Non-Mobilizable | 3916 | - | - |
| scaffold_12628 | 35 | scaffold_9153 | 37 | Non-Mobilizable | 4388 | - | - |
| scaffold_35161 | 29 | scaffold_25359 | 47 | Non-Mobilizable | 2218 | - | - |
| scaffold_12673 | 28 | scaffold_9299 | 17 | Non-Mobilizable | 4335 | - | Toxin-coding gene *yoeB*,<br>Antitoxin-coding gene *yefM* |
| scaffold_130 | 19 | scaffold_49 | 88 | Non-Mobilizable | 94434 | Metal Resistance (Cobalt Zinc, Cadmium, Copper, Arsenic), Antibiotic Resistance (Fosfomycin, Spectinomycin), Metal uptake (Potassium uptake) | - |
| scaffold_3845 | 17 | scaffold_2793 | 17 | Conjugative | 10328 | - | - |
| scaffold_34293 | 10 | scaffold_24781 | 298 | Non-Mobilizable | 2253 | - | - |
| scaffold_23147 | 8 | scaffold_16750 | 191 | Conjugative | 2916 | - | - |
| scaffold_31933 | 7 | scaffold_23032 | 12 | Non-Mobilizable | 2363 | - | - |
| scaffold_27134 | 6 | scaffold_19474 | 325 | Conjugative | 2632 | - | - |

**Fig 3.**
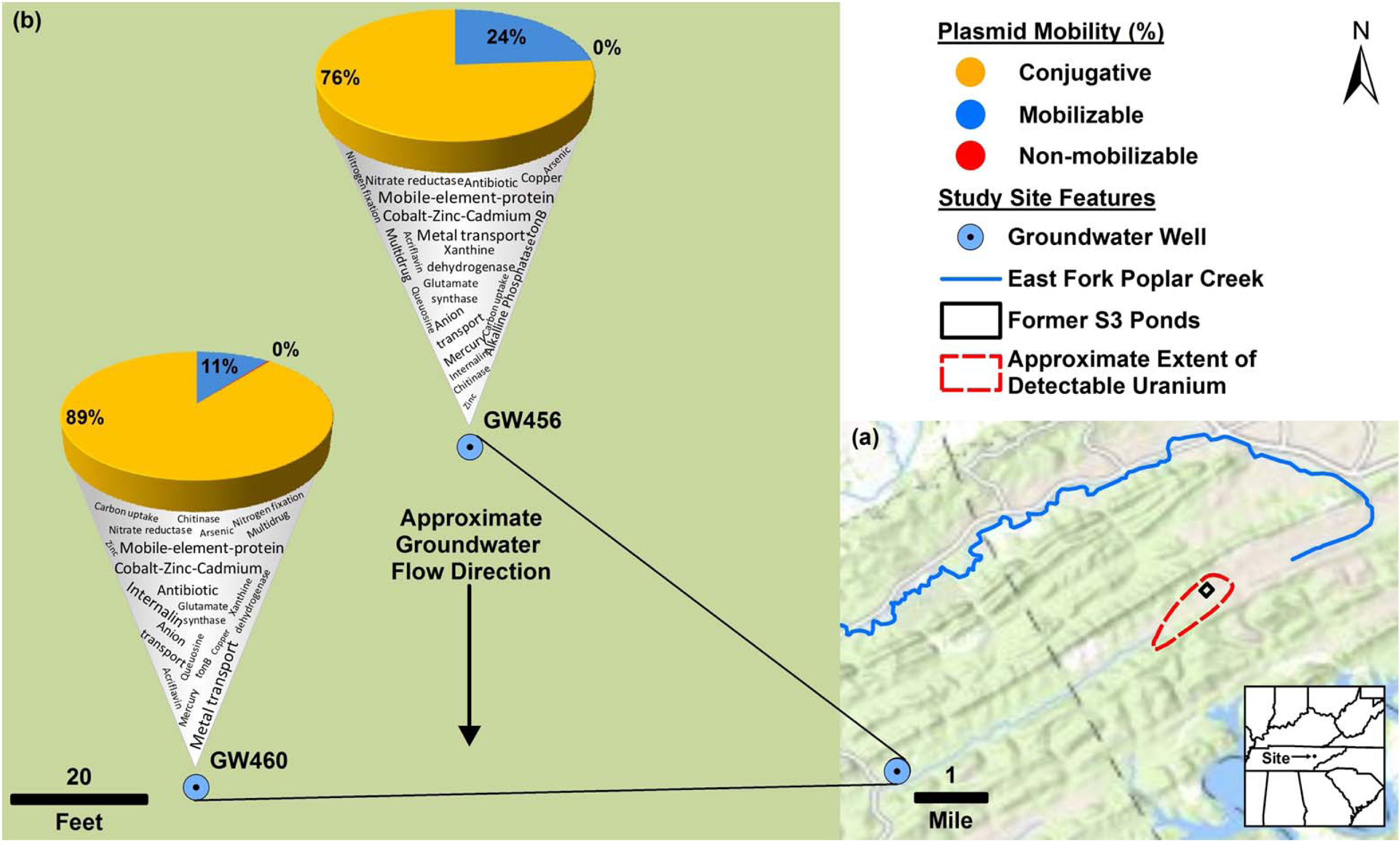
Plan-view maps of the OR-FRC, showing large- and small-scale features, (a) and (b), respectively. Approximate extent of detectable uranium in (a) based on Fig. 1 from Smith et al., (2015). Mercury has been detected at East Fork Poplar Creek (Brooks and Southwort, 2011). b) displays the mobility of “circular_scaffolds” (as a percentage of total circular plasmids in that well) along with a word cloud denoting the Kbase annotated functions encoded on the “circular_scaffolds”. Approximate groundwater flow direction in based on Fig. 7 from Moline et al., (1998). The base map was provided by Environmental Systems Research Institute (ESRI).

Next, we compared the “circular_scaffolds” and the “all_scaffolds” sequences to specific databases focused on plasmid, antibiotic resistance and metal resistance genes. The ACLAME plasmid database contains genes from more than a thousand plasmids (Lima-Mendez *et al.*, 2010). The wells GW456 and GW460 had 55.3 and 43.5 % of “all_scaffolds” and 80.6 and 70.6 % of “circular_scaffolds” with hits against the ACLAME plasmid database. This indicates that plasmid sequences obtained are similar to those reported to be plasmid-associated. The most abundant gene source based on ACLAME were hypothetical proteins, and ABC transporter related genes (Supplementary Information 7).

Analysis of “all_scaffolds” for known antibiotic resistance genes revealed the top categories in both wells to be aminocoumarin resistance and elfamycin resistance when queried against CARD database, and bacitracin resistance when queried against ARDB (Supplementary Information 8).

Also, chloramphenicol resistance genes were highly abundant in well GW460. A search using the “circular_scaffolds” against ARDB yielded no hits, whereas on CARD it yielded 2 (rifampicin and elfamycin resistance) and 0 hits in GW456 and GW460, respectively.

Toxin-antitoxin systems are known to be important in plasmid maintenance (Jaffé *et al.*, 1985; Gerdes *et al.*, 1986). A search against the toxin-antitoxin database revealed that less than 1 % of “all_scaffolds” encoded both toxin and antitoxin genes. However, a higher percentage of “circular_scaffolds” (19.1 and 7.7 % in wells GW456 and GW460) encoded both toxin and antitoxin genes (Supplementary Information 9). Interestingly, certain circular plasmids also contained CRISPR repeats, spacer and Cas proteins. Certain scaffolds such as scaffold_295 from the well GW456 contained CRISPR associated Cas1-4 along with Doc toxin, and Phd antitoxin protein known to be associated with plasmids (Gerdes *et al.*, 2005; Liu *et al.*, 2008), hinting that viral resistance might also be a plasmid encoded trait.

## Plasmid-borne metal resistance genes are prevalent in the pristine wells

We found our most informative ecologically relevant results while evaluating heavy metal resistance genes in the sequenced scaffolds. Heavy metal resistance genes are one of the most frequently found phenotypic modules carried by bacterial plasmids (Silver, 1996). We subjected “all_scaffolds” from both wells to the BacMet database (Pal *et al.*, 2014) to obtain gene calls by comparison with predicted and experimentally confirmed antibacterial biocide- and metal-resistance genes (Supplementary Information 10). Out of the 553 genes in the predicted category, hits were found against 212 and 221 genes in the wells GW456 and GW460, respectively. Of these the top most abundant genes were those annotated to belong to *copR* (copper-responsive regulators), *arsB* (arsenic specific transporter), *merA* (mercuric ion reductase) in GW456, and *ruvB* (Zinc metalloprotease), *mexT* (regulator of multidrug efflux pump), and *copR* in GW460. As observed earlier at the OR-FRC (Hemme *et al.*, 2016), the Cd2+/Zn2+/Co2+ (*czc*) efflux genes were also highly abundant. Proteobacteria was the most abundant gene source for the top metal resistance encoding genes (Fig 4). Despite the absence of any currently elevated concentrations of heavy metals, a previous study noted high abundance of metal resistance genes in the pristine wells at OR-FRC (Hemme *et al.*, 2015), and hypothesized that they might be present on a plasmid. This study confirms that several metal resistance genes are indeed encoded on plasmids.

**Fig 4.**
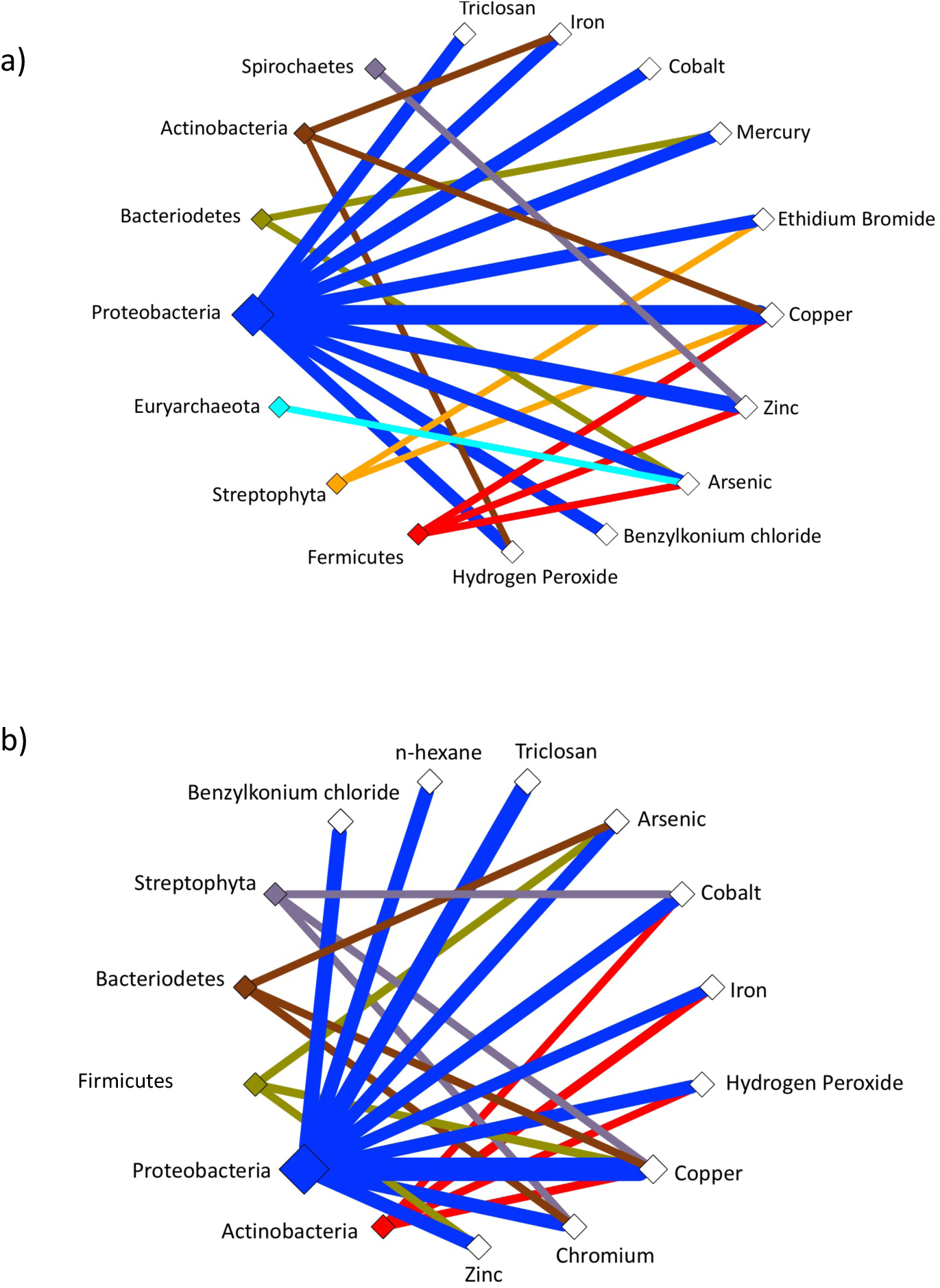
Metal resistance genes of Proteobacterial origin are abundant in the pristine wells at the Oakridge site. Cytoscape was used to generate a network of predicted bacterial phyla coding for resistance genes as per the BacMet database. Edge thickness indicates the number of genes encoded in a) GW456 and b) GW460.

An example is the 170 kb circular plasmid-scaffold_67, that carries several metal resistance genes such as those involved in Cobalt, Zinc and Cadmium resistance (*czcABCD*) along with Copper resistance (*copG*). This is a mobilizable plasmid, carrying genes *traIDALEKBCWUNFHG* along with *trbC.* It also encodes a response regulator *phoQ.* The entire plasmid depicts 94 % identity to *Sphingobium baderi* DE-13 plasmid pDE1 (10 % query coverage). Interestingly, the strain *Sphingobium baderi* DE-13 was isolated from an activated sludge from an herbicide-manufacturing factory in Kunshan, China (Li *et al.*, 2013).

One of the most striking observations of our study is that the most abundant circular plasmid common to both wells encodes mercury resistance. The circular scaffold_5343 displayed the highest coverage (869x coverage) and is hence the most abundant plasmid in GW456 (Fig 5). The circular scaffold_10032 (617x coverage) from GW460 displayed 100 % identity (98 % query coverage) to scaffold_5343, indicating this plasmid was highly abundant in both wells. This plasmid codes for genes annotated to be involved in mercury uptake and resistance along with plasmid mobilization and replication genes. Most of the genes on this plasmid had homologs in the genus *Paracoccus.* The abundance of the genus *Paracoccus* was merely 0.16 % based on 16S rRNA abundance in the well GW456, perhaps indicating that the plasmid might have consequently horizontally transferred into other hosts and/or is maintained in the original host in high copy numbers.

**Fig 5.**
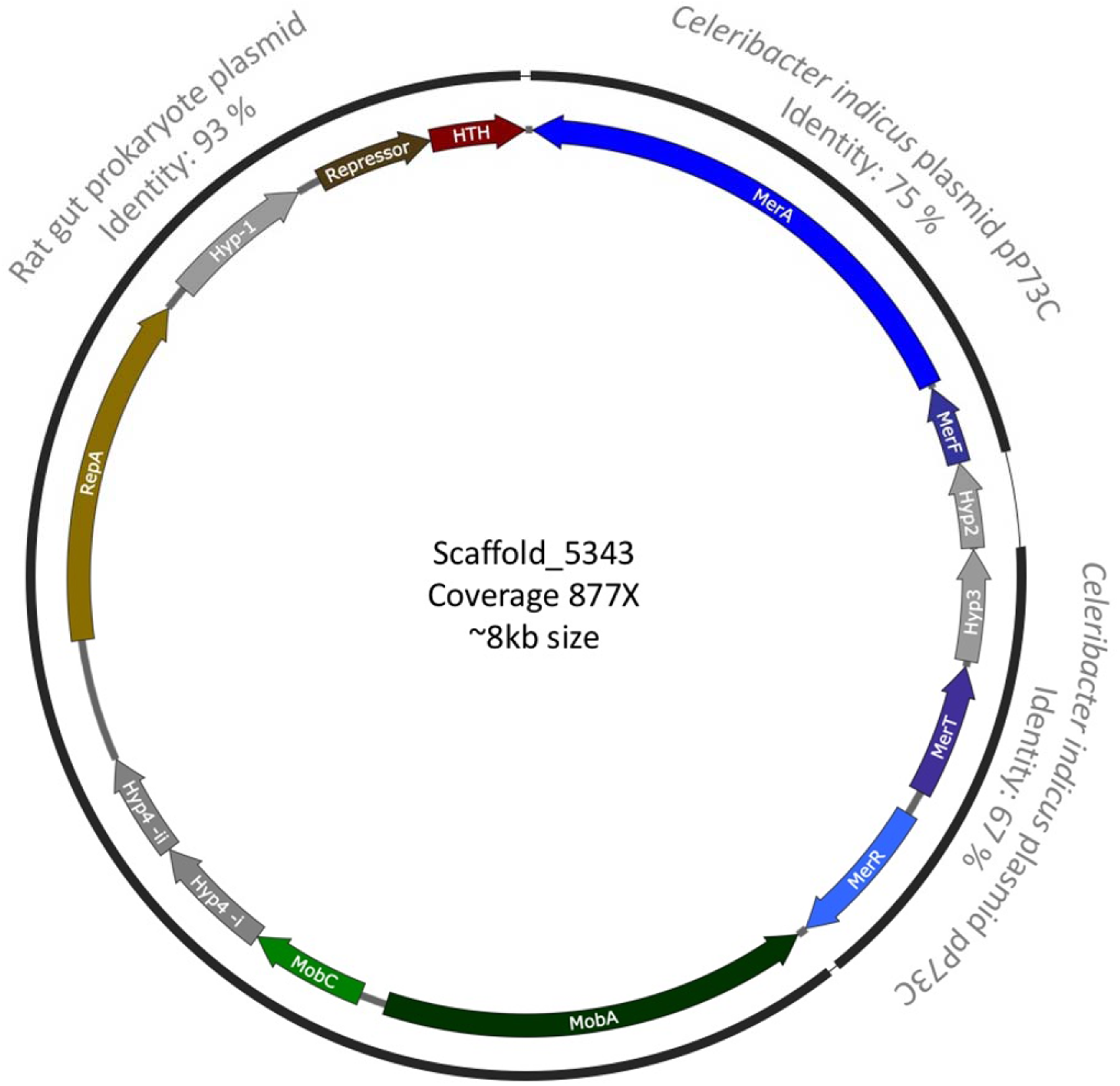
Plasmid map of the most abundant plasmid in well GW456. Genes encode the following proteins MerA: Mercuric ion reductase, MerF: Mercuric ion uptake protein, Hyp: Hypothetical protein, MerT: Mercuric transport protein, MerR: Regulator of Mercury resistance genes, MobA: Mobilization protein A, MobC: Mobilization protein C, RepA: Plasmid replication protein, HTH: Helix-turn-helix domain protein, Repressor: Cytotoxic repressor of toxin-antitoxin. The plasmid was randomly cut by IDBA-UD in the hypothetical protein 4, and hence the 2 parts of the same protein are depicted in the figure. The black lines indicate the plasmid can be broken into different modules that show similarity to other previously reported plasmids (the closest NCBI blast hit with more than 92 % query coverage depicted in grey).

We also performed a nucleotide blast on the whole plasmid sequence of the circular scaffold_5343 which shows that this plasmid can be broken into three modules. The first module spans from *mobA* to the helix-turn-helix containing protein and exhibits homology to a rat gut plasmid GenBank: LN852881.1 (total size 12.9 kb). The original rat gut plasmid codes for certain hypothetical proteins in addition to the *ccdA/ccdB* Type II toxin-antitoxin system genes. The second and third modules encode mercury resistance genes and depict homology to the native plasmid pP73c (total size 122 kb) in *Celeribacter indicus* P73^T^. The P73^T^ strain was isolated from a deep sea sediment in the Indian Ocean (Cao *et al.*, 2015).

Since the ground water samples show no presence of mercury, this study points us to an interesting question – why have plasmids with metal resistance genes persisted in pristine environments? For the plasmid to be maintained, it must either confer a selective advantage to the host or replicate faster than the hosts. If not, they are bound for extinction (la Cruz and Garcillán-Barcia, 2014). Another study (Millan *et al.*, 2014) demonstrated that plasmid persistence could be attributed to compensatory adaption, along with brief periods of positive selection, which might be the most plausible explanation for the persistence of metal resistance gene(s) on plasmids in the pristine wells. Even then, the persistence over long periods of time might be linked to benefits of encoding gene on a plasmid rather than the chromosome, such as obtaining higher levels of expression. It is also possible that most plasmids observed in our study can be tracked back to the few common bacteria present in both wells. Our study suggests that the microbial community in pristine wells is likely robust in tolerating low stresses and possesses a latent ability to swiftly adapt to changes in the environmental stress levels.

## Conclusion

This is the first study to selectively isolate and analyze the plasmid population from a unique environment such as the OR-FRC. Given the 1) low cell density of the pristine wells, 2) absence of contaminants coupled with 3) the burden associated with carrying plasmids, it was surprising to find a rich plasmidome community in these wells. Technically, it is difficult to avoid biasing plasmid isolation in favor of smaller plasmids. Previous studies on plasmidome analysis have relied on commercial kits (better at isolating small plasmids) (Jørgensen *et al.*, 2014b; Zhang *et al.*, 2011) or pooled DNA from various plasmid isolation methods (Kav *et al.*, 2013) along with phi29 amplification at 30 °C. We optimized the plasmid isolation and amplification to be able to better detect large plasmids present in low copy numbers in the environment and were successful in obtaining large complete circular plasmid units.

Even though the taxonomic distribution of bacteria varies from well to well when categorized based on 16S sequences, the variation is much less when evaluated based on plasmidome, and may have ecological significance in the role of plasmids in maintaining key latent functionalities. The presence of metal resistance genes, specifically of highly conserved mercury resistance genes on the most abundant plasmids, is noteworthy due to the lack of mercury or other heavy metal contamination in wells GW460 and GW456. Mercury is one of the major contaminants at the OR-FRC (He *et al.*, 2010), and it was suggested that mercury resistance genes were horizontally transferred in this environment (Martinez *et al.*, 2006). The plasmidome analysis of this site provides evidence that plasmid-mediated HGT plays a crucial role in driving the evolution of a groundwater microbial community in response to contamination. Our specific observations are unique to the OR-FRC microbiomes, but the methods to examine plasmid DNA and their significance are generalizable to all microbial communities.

## Acknowledgements

This work was part of the ENIGMA- Ecosystems and Networks Integrated with Genes and Molecular Assemblies (http://enigma.lbl.gov). a Scientific Focus Area Program at Lawrence Berkeley National Laboratory and is supported by the U.S. Department of Energy, Office of Science, Office of Biological & Environmental Research under contract number DE-AC02-05CH11231 between Lawrence Berkeley National Laboratory and the U. S. Department of Energy. The United States Government retains and the publisher, by accepting the article for publication, acknowledges that the United States Government retains a non-exclusive, paid-up, irrevocable, world-wide license to publish or reproduce the published form of this manuscript, or allow others to do so, for United States Government purposes.

